# A Novel Mechanism for Tauopathy in Progressive Multiple Sclerosis: Excitotoxic Misplacement of a Mitochondrial Anchor into Dendrites Driven by Tau-hyperphosphorylation

**DOI:** 10.1101/2025.09.05.674541

**Authors:** Deepali Mathur, Charlie Zhang, Dinesh C Joshi, Jonathan E. Ploski, Shing Y Chiu

## Abstract

On April 2^nd^, 2025, the FDA approved a Fast Track Designation for Biogen to use Antisense Oligonucleotides (ASO) to treat tauopathy in clinical trials for Alzheimer’s Disease (AD) to meet an unmet medical need [1]. For Multiple Sclerosis (MS), there is a similar unmet medical need regarding tauopathy when MS transitions into the late, or Progressive MS that is currently incurable. AD and MS share commonality: there is comorbidity between AD and MS [2], and the recent awareness that progressive MS may be considered a secondary tauopathy [3]. This study lays the basic science foundation for a future repurposing of ASO tauopathy therapy from AD to MS. The central hypothesis is that in Progressive MS, tauopathy is not a passive *bystander* but an active contributor to synaptic degeneration through a novel toxic target known as **<u>DSI</u>** (**<u>D</u>**endritic **<u>S</u>**yntaphilin **I**ntrusion) discovered in our laboratory. In this hypothesis, the excitotoxic N-methyl-D-aspartate receptor (NMDAR) GluN2B activates Tau hyperphosphorylation (p-Tau), leading to the mislocalization or intrusion of a mitochondrial anchor SNPH into neuronal dendrites (DSI). This causes mitochondrial damage and subsequent synapse/dendrite disintegration. In support of this hypothesis that tauopathy is a key driver of DSI, we demonstrated using primary neuronal cultures that inhibitors of p-Tau kinases and Tau-KO both completely abolish DSI. We propose that a therapy for Progressive MS is repurpose the existing FDA-approved ASO Tau knockdown therapy from AD to treat MS.

**Funding:** NIH R01 MSN233781 to SY Chiu

## 1. Introduction

Even though Multiple Sclerosis (MS) is known classically as a white matter axon demyelination disease, gray matter consisting of cell bodies and dendrites of neurons is an important pathology in MS independent from white matter demyelination [4-10]. Indeed, gray matter damage is a major driver of disability in progressive MS [11-13]. Intriguingly, progressive MS bears similarity to Alzheimer’s Disease (AD) in exhibiting characters of a tauopathy disease. While it is well established that hyperphosphorylated tau (p-Tau) is a pathological hallmark of Alzheimer’s disease (AD), accumulating evidence [14] indicates that altered tau phosphorylation may be driving multiple sclerosis (MS) pathology. For example, tau phosphorylation increases in a key mouse model of MS [15]. Notably, in the untreatable progressive phase of MS, cerebrospinal fluid (CSF) p-Tau increases over time, reaching the highest levels in the late phase of primary progressive MS (PPMS)[16]. The prominence of p-Tau in late-phase MS suggests that progressive MS may be considered a secondary tauopathy [3]. An important therapeutic issue regarding tauopathy in progressive MS is whether phosphorylated tau is just a ‘bystander’ of disease progression or whether it actively contributes to neurodegeneration in MS [14].

In this study, we propose that in progressive MS, p-Tau is not a bystander but an active driver of neurodegeneration via a novel excitotoxic target, **<u>DSI</u>**, recently discovered in our laboratory in a mouse model of progressive MS [17]. **<u>DSI</u>** stands for **<u>D</u>**endritic **<u>S</u>**yntaphilin **I**ntrusion. Syntaphilin (SNPH) is a mitochondrial anchoring protein normally expressed in axons. In progressive MS rodent models, we discovered an abnormal mislocalization of SNPH into dendrites (**<u>DSI</u>**), causing excitotoxic damage to mitochondria and neurons [17]. Our hypothesis that p-Tau drives the excitotoxic DSI to damage progressive MS carries enormous therapeutic implications: Existing FDA-approved tauopathy therapy for AD can be expedited and repurposed to treat MS.

Our hypothesis that Tau drives DSI is summarized in Fig. 1 below.

**Fig. 1.**
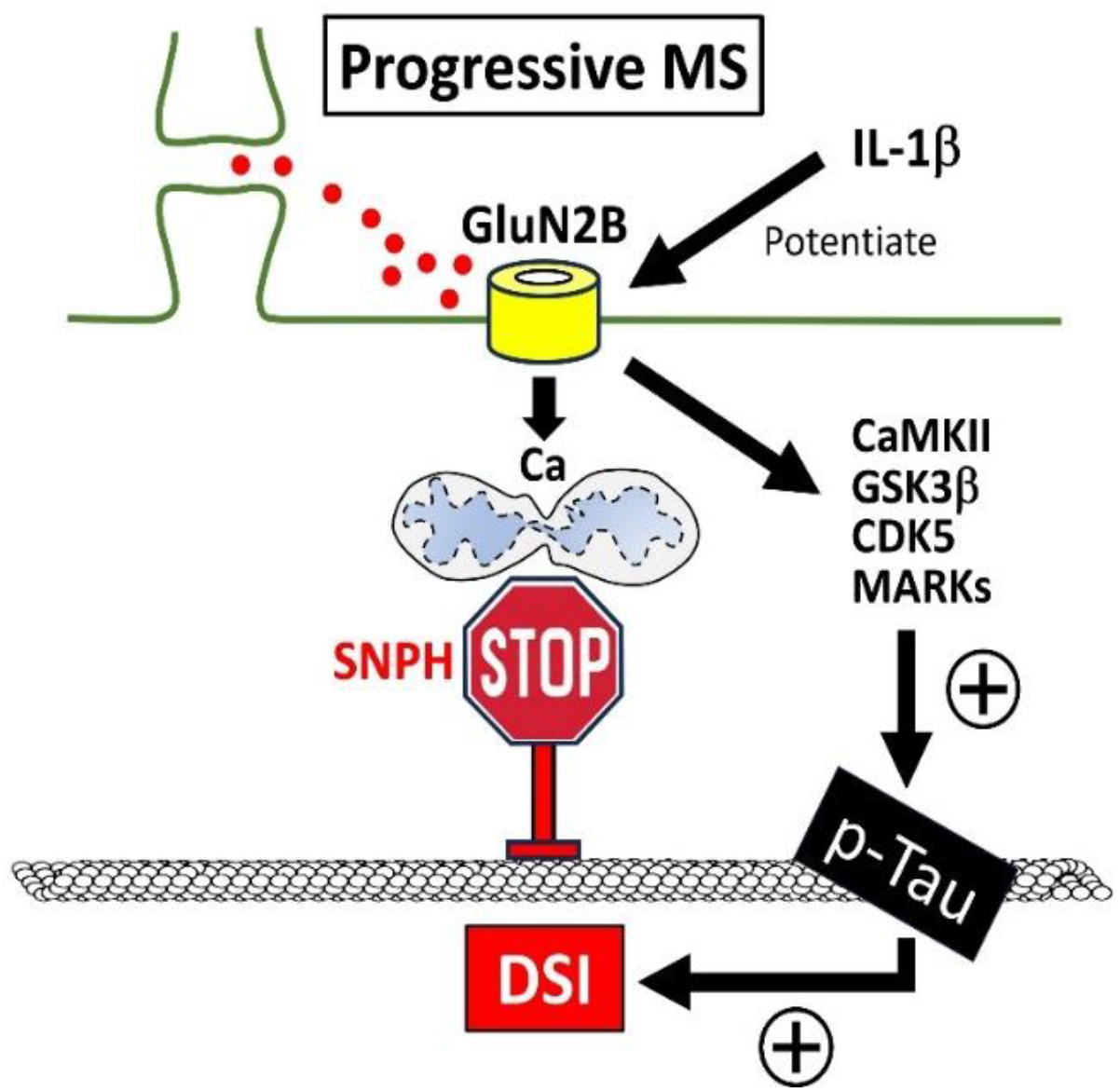
Hypothesis on Pathologic Linkage between Tau and DSI in Progressive MS. a) Gray matter dendritic damage is a major driver of pathology in Progressive MS [11-13]. b) In Progressive MS, extrasynaptic GluN2B is potentiated by pro-inflammatory cytokines IL-1β and glutamate spill-over from synapse. c) Hyperactivation of GluN2B activates various kinases CaMKII, GSK3β, CDK5 and MARKs to trigger hyperphosphorylation of Tau (p-Tau). d) p-Tau drives DSI to immobilize mitochondria in dendrites close to GluN2B. e) GluN2B generates excitotoxic Ca^2+^ to overload mitochondria leading to mitochondrial fragmentation and dendrite degeneration. f) Knockdown of Tau by Tau-KO or Tau ASO will abolish DSI, prevent gray matter damage and protect progressive MS.

The goal is to provide key supporting data for the tauopathy pathway in the hypothesis in Fig. 1: GluN2B triggers p-Tau to drive the excitotoxic DSI.

## 2. Materials and Methods

### 2.1. Experimental Animals

Mice of either sex were used in this study. All experiments involving animals were performed in accordance with University of Wisconsin–Madison Research Animal Resources and Compliance. All animal usage and protocols were reviewed by the Animal Care and Usage Committee and approved by University of Wisconsin–Madison Research Animal Resources Center. Wt NOD mice were obtained from Charles River laboratory and NOD-SNPH-KO mice were generated at Genome Editing and Animal Models Facility at UW Madison. Homozygous C57BL6 WT and Tau-KO mice were obtained from the Jackson Laboratory.

### 2.2. Induction of EAE in NOD and NOD-SNPH-KO

NOD mice were immunized by subcutaneous injection of 200µg (100µg in each flank) of MOG35-55 peptide (Genemed Synthesis) in complete Freund’s adjuvant (Sigma-Aldrich) containing 5 mg/ml mycobacterium tuberculosis H37Ra (BD Biosciences). Mice were also injected with 200 ng of pertussis toxin on day 0 and 2 by tail vein or intraperitoneal injection. Following the injection, mice were scored for clinical symptoms on a 0 –5 scale as follows: 0-no clinical symptoms,1-limp/flaccid tail, 2-hindlimb weakness with incomplete paralysis, 3-complete hindlimb paralysis, 4-paraplegia, and 5-moribund or dead.

### 2.3. Immunofluorescence and image acquisition

For immunofluorescence analysis, tissue from cerebellum were processed for paraffin embedded sections at TRIP pathology core facility at UW Madison. The glass slide containing sections were washed with xylene and hydrated through a series of alcohol and water. The slides were boiled in antigen unmasking solution (Vector laboratories) for 20min and washed with PBS 3 times followed by incubation with blocking solution for 1hr in 0.3% Triton X-100, 10% normal goat serum prepared in 1X PBS. Tissue sections were then incubated with primary antibodies SNPH (Abcam, 1:250), Syt2 (ABCam, 1:200), Calbindin (ABCam, 1:250) overnight at 4-degree C. Following day sections were washed with 1X PBS (5 times, 5min each) and incubated with Alexa fluor labelled secondar antibodies for 1hr and washed with 1X PBS (5 times, 5min each) before mounting with UltraCruz Aqueous Mounting Medium (Santa Cruz Biotechnology). Fluorescence images were acquired using Nikon A1 confocal microscope (UW optical imaging core facility) with 10x, 20x and 60x objectives at 512X512 and 1024X1024 pixel resolution.

### 2.4. Culture neurons from wild type and Tau-KO mice

Mouse primary hippocampal neurons were isolated from newborn postnatal day 0 (P0) C57BL6 WT and Tau-KO (obtained from the Jackson Laboratory). Briefly, P0 mice were decapitated, and hippocampi were isolated in ice-cold high glucose DMEM (HGDMEM) plus 10% fetal bovine serum (FBS). The hippocampi were rinsed twice with ice-cold HGDMEM and incubated with Papain (2 mg/ml prepared in HGDMEM) for 30 min at 5% CO_2_ at 37°C. After digestion in papain, DNase was added (2.5 mg/ml prepared in HGDMEM) for 30 s followed by the addition of 10% FBS to inactivate DNase. The digested tissue was gently triturated by pipetting and filtered through a 100 µm sterile cell strainer and spun at 800 rpm for 10 min at room temperature (RT). Cell pellets were resuspended in a 3:1 ratio of neurobasal-B27 and HGDMEM (with 10% fetal bovine serum) and plated on poly-D-lysine-coated (0.2 mg/ml) coverslips at a density of 1.1 × 105 cells/ml. The plated neurons were switched to neurobasal B27 medium after 5 h and were treated with 1 μM Ara-C after 48 h of plating for 48 h to inhibit the growth of non-neuronal cells in the culture. Cells were then maintained in neurobasal B27 medium with one-third of the media changed every 48 h. DSI experiments were performed on neurons cultured for 9–12 d.

### 2.5. DSI imaging analysis

Dendritic SNPH is measured by orthogonal analysis. Cultured neurons were stained with MAP2 to identify dendrites and SNPH to identify DSI. To verify that SNPH is inside dendrites, orthogonal images at three different angles were obtained using NIS-Elements software (Nikon) with a 60× oil lens under a Nikon A1RS Confocal Microscope at the University of Wisconsin–Madison Optical Imaging Core to confirm that SNPH immunoreactivity originates from inside the MAP2-positive dendrites. To quantify DSI, we first open the 3D images with NIS-Elements to identify if yellow puncta (SNPH) are inside red dendrites (green puncta, SNPH, are outside dendrites) by orthogonal analysis under 1000% magnification. At the same time, we open the same image with ImageJ software on another computer to draw contours of all SNPH immunoreactivity (yellow puncta) within the total MAP2 area (red areas) within the fixed frame. This fixed frame is standardized and applied to all samples. Both areas of yellow and red are measured and recorded by ImageJ and saved in Microsoft Excel. Finally, DSI is computed as the total SNPH area divided by the total MAP2 area for a particular dendritic segment under analysis with Excel. Analysis was performed by one of us who was blinded to the experimental conditions.

### 2.6. Statistics

Data are presented as the mean ± SEM. Statistical differences in measured variables between the experimental and control groups were assessed by Student’s unpaired *t* test. One-way Anova followed by Post Hoc Tukey HSD (beta) test was applied when comparison was made between more than two variables and *p* < 0.05 was considered statistically significant.

## 3. Results

### 3.1 DSI is present in two progressive MS mouse models (PPMS and SPMS)

In humans, the entire spectrum of MS is encompassed by two disease phenotypes: Relapsing remitting MS (RRMS), majority of which eventually transforms into secondary progressive MS (SPMS) and primary progressive MS (PPMS). It is estimated that 85% of patients are primarily affected with RRMS out of which roughly 65% are converted into SPMS over a period of time. PPMS constitutes 15% of the entire MS population. PPMS and SPMS have distinct disease phenotypes. In PPMS, patients start with a steadily increasing neurologic dysfunction without periods of remission or relapses. In contrast, SPMS patients start with a relapsing-remitting phase (RRMS) before transitioning to the secondary progressive phase (SPMS) of steadily increasing disease progression. In 2019 [17], we first discovered DSI in a PPMS mouse model (*Shiverer* mice). We now extended the presence of DSI to a SPMS mouse model. For the SPMS model, we used the Experimental Autoimmune Encephalomyelitis (EAE) mouse in the NOD (<u>N</u>on-<u>O</u>bese-<u>D</u>iabetic) background [18]. The EAE/NOD mice (EAE induced in 8-week-old mice) exhibit an early RRMS phase that transitions to a prominent SPMS phase after ∼70 days and beyond [18].

Fig. 2 shows DSI analysis in the dendrites of Purkinje cells in the cerebellum from NOD/EAE at 180 days post-immunization, well into the progressive phase. Dendrites were labelled with calbindin.

**Fig. 2.**
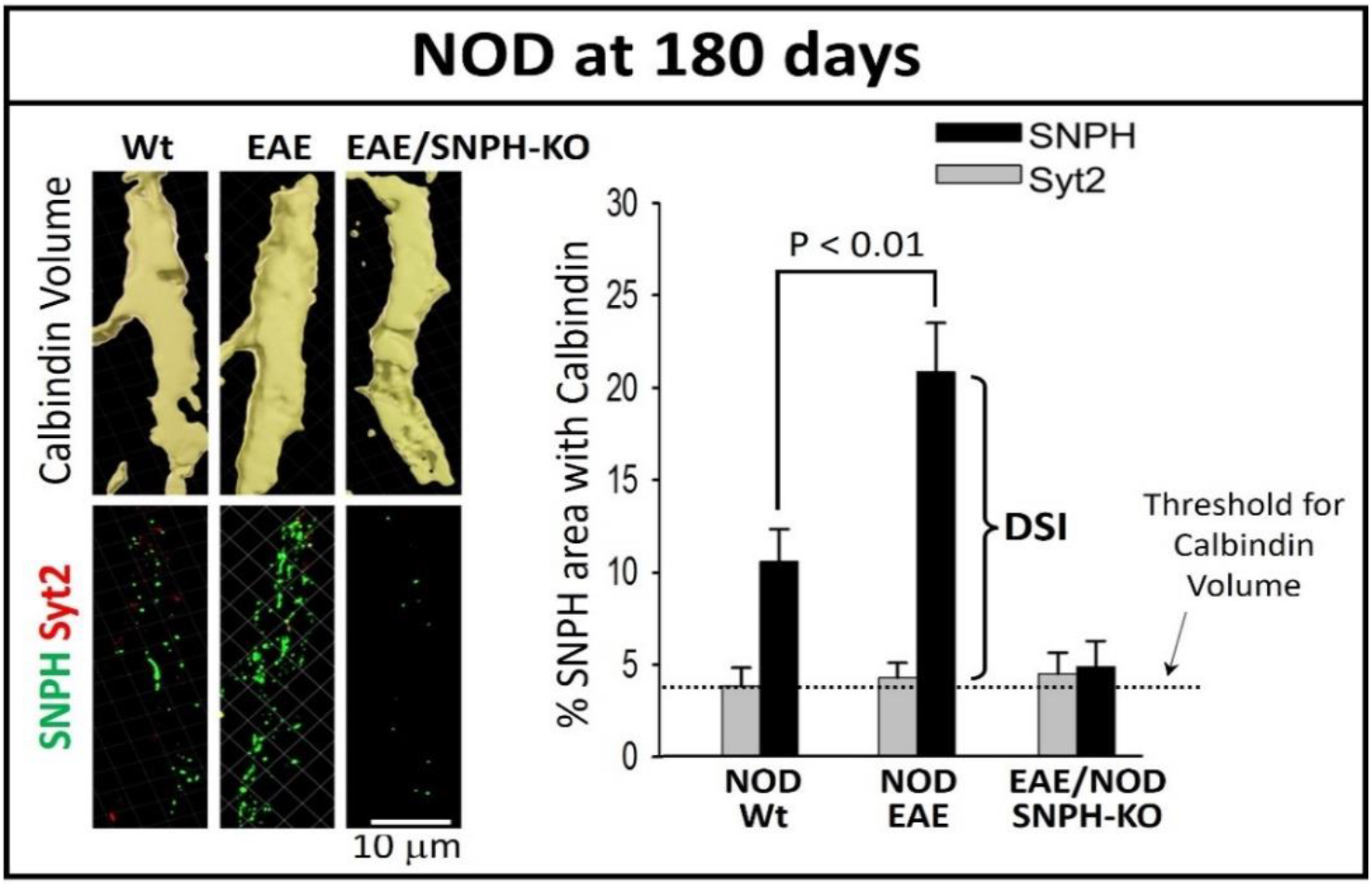
DSI in Purkinje cell dendrites in cerebellum in SPMS model (EAE/NOD mouse) Cerebellar sections of Wt, EAE, EAE/SNPH-KO in NOD mice at 6 months post-immunization were labeled with SNPH, Syt2, and Calbindin antibodies. After deconvolution, dendritic regions were highlighted by adding a surface mask onto the threshold Calbindin channel using Imaris software. The SNPH and Syt2 fluorescence signals outside the dendrite were then removed to quantify the percentage of SNPH and Syt2 within the Calbindin volume (N=8 dendrites from 3 mice). Extensive DSI is present in EAE but abolished in EAE/SNPH-KO. For details of methods, consult our publication (Fig. 1 of Joshi et al., 2019 [17]).

To demonstrate rigor, we measured DSI using a method called calbindin volume reconstruction (based on our work [17]). Briefly, we reconstructed the calbindin volume in Purkinje cells using the presynaptic marker Syt2 to exclude axonal contributions to the dendritic volume. The reconstructed dendritic calbindin volume using Syt2 as the exclusion marker is shown in Fig. 2 for NOD (upper images). The bottom panels show double staining for **SNPH** (**Green**) and the presynaptic marker **Syt2** (**Red**) of the reconstructed calbindin volume showing successful calbindin volume reconstruction with little or no **Syt2** (**Red**) contributions from axonal volumes. Fig. 2 shows extensive DSI present in the dendritic calbindin volume in EAE/NOD at 180 days compared to control NOD/Wt without EAE.

To decisively confirm that DSI is genuine, we used CRISPR gene editing at the University of Wisconsin Madison Core Facility to generate SNPH-KO in the EAE/NOD mice to demonstrate that SNPH-KO completely eliminate the DSI signal, demonstrating that our DSI signal has no non-specific artifacts.

Collectively, we established that DSI is present in the progressive phase of two MS mouse models, both in the *Shiverer* mouse model for PPMS [17], and the EAE/NOD mouse model for SPMS (Fig. 2).

### 3.2 NMDA differentially activates GluN2B to trigger DSI

Having discovered DSI in Progressive MS models, we next explored MS-relevant pathological signals that trigger DSI using cultured neurons. Since glutamate excitotoxicity is considered to be a significant factor in MS lesions [19], we recently published that in cultured hippocampal neurons, NMDA triggers DSI [17, 20]. In the current study, we examined the NMDAR subtypes involved in triggering DSI. The NMDAR has two subtypes, GluN2A and GluN2B, playing opposing roles in excitotoxicity: GluN2A is pro-survival and GluN2B is pro-excitotoxic. In Fig. 3, we found that inhibiting GluN2B with RO25 completely prevents NMDA from triggering DSI (Fig. 3C, D), thereby identifying GluN2B as the key NMDAR subtype driving DSI.

**Fig. 3.**
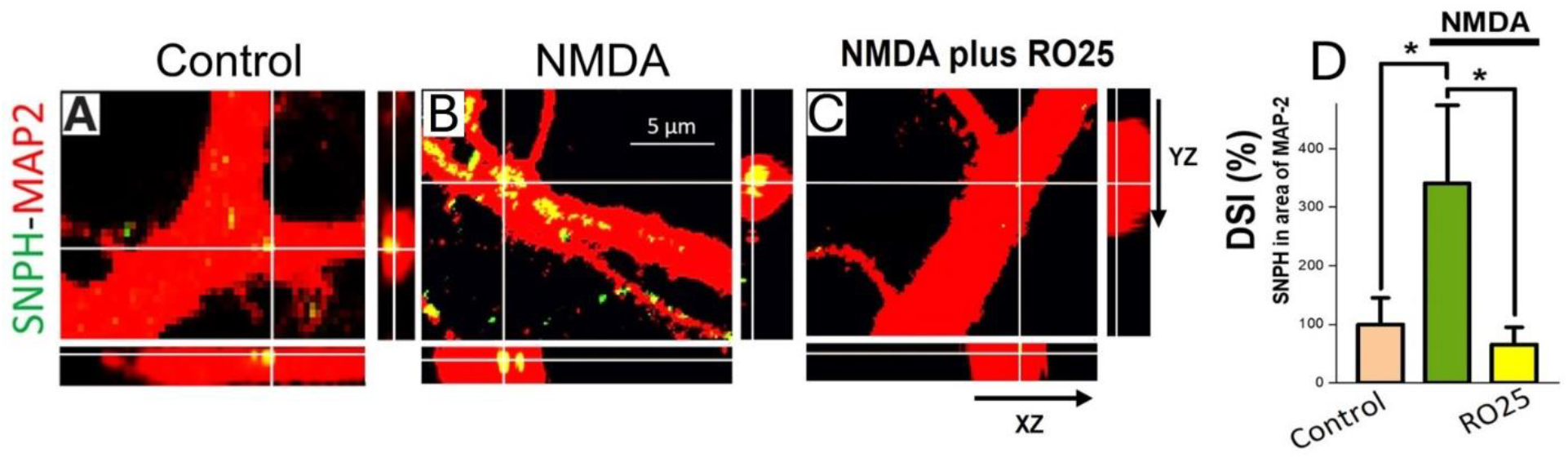
NMDAR-GluN2B trigger DSI in cultured hippocampal neurons. Neurons treated NMDA (10μM, 24 h) then double stained for SNPH and MAP2 to detect DSI. For DSI, the SNPH puncta are inside dendrites and turn yellow in the merged SNPH-MAP2 images (A, B) as verified by orthogonal analysis (bottom and right images). (C) Selective inhibition of GluN2B with RO25-6981 (10 μM) completely blocked DSI triggered by NMDA, identifying GluN2B as the key NMDAR subtype driving DSI. Cells were pre-treated with inhibitors for 30 min before NMDA application. (D) - Quantification of the inhibitory effects RO25 (10μM) on DSI induced by NMDA. N=30 dendrites for each group. (* p<0.05, ***p<0.001).

We further corroborate the pivotal role of GluN2B in driving DSI by contrasting it with GluN2A using viral transduction.

Fig. 4 shows the effect on ambient DSI by transducing cultured neurons with GluN2A and GluN2B encoding viruses. GluN2A virus has no effect on DSI (A, C). In striking contrast, GluN2B virus (B, D) dramatically increased DSI under comparable viral transduction measured by FLAG immunocytochemistry (E). GluN2B virus dramatically sensitizes neurons to DSI, where basal release of glutamate by neurons in culture is sufficient to trigger ambient DSI. In Progressive MS, two factors further augment the pathological effect of GluN2B according to our hypothesis in Fig. 1. First, low levels of pro-inflammatory cytokine, IL-1β (like that found in MS patients), have been shown to potentiate the activity of extrasynaptic NMDA receptors, including GluN2B [21]. Second, dysfunctional glutamate uptake in MS can cause glutamate spill-over from the synapse to activate the extra-synaptic GluN2B.

**Fig. 4.**
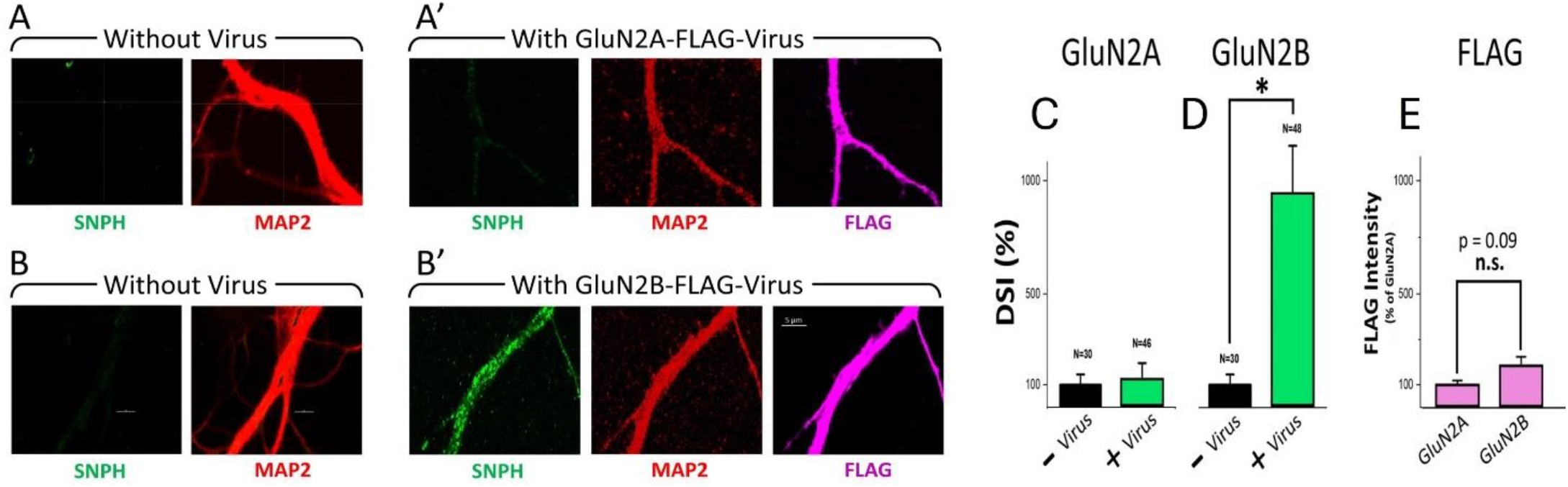
DSI is driven by GluN2B and not GluN2A. Cultured hippocampal neurons were treated with 200 µl of Lenti-CMV Flag-GluN2A Wt and Lenti-CMV Flag-GluN2B Wt virus on first day of culture and permeabilized and fixed with 4% PFA on sixth day for SNPH, MAP2 and FLAG immunohistochemistry. Ambient DSI unaffected by GluN2A virus (A, C) but dramatically increased by GluN2B virus (B, D). Dendrites N (30 non-virus, 46 GluN2A, 48 for GluN2B). Unpaired t test * p<0.05 between control and GluN2B. (E) No statistical difference (n.s. p=0.09) found between FLAG intensities of GluN2A and GluN2B expression. FLAG intensity measurement in dendrites treated with GluN2A-FLAG or GluN2B-FLAG viruses. FLAG measured in fixed, permeabilized neurons by ImageJ software in dendrites of GluN2A (N=30) and GluN2B (N=28); Unpaired t test applied.

Collectively, the extra-synaptic GluN2B is a strong candidate for exacerbating the toxic DSI to damage Progressive MS.

### 3.3. GluN2B activates p-Tau to drive DSI

The discovery that GluN2B plays a pivotal role in driving DSI is crucial for our p-Tau hypothesis. It has already been demonstrated that GluN2B activates CaMKII to trigger p-Tau, and inhibiting CaMKII completely blocks GluN2B from triggering p-Tau. We therefore examined if preventing p-Tau by blocking CaMKII with KN-93 also blocks GluN2B from triggering DSI.

Excitingly, Fig. 5 shows that blocking CaMKII with KN93 prevents NMDA from driving DSI (Fig. 5C, G). Besides CaMKII, we explored the roles of three other kinases known to promote p-Tau. The first is GSK3β that is known to be elevated in brain regions of patients with progressive MS [23]. The second is CDK5 that is activated in MS [24] and the EAE model [8]. The third is MARKs, a key kinase for tau phosphorylation.

**Fig. 5.**
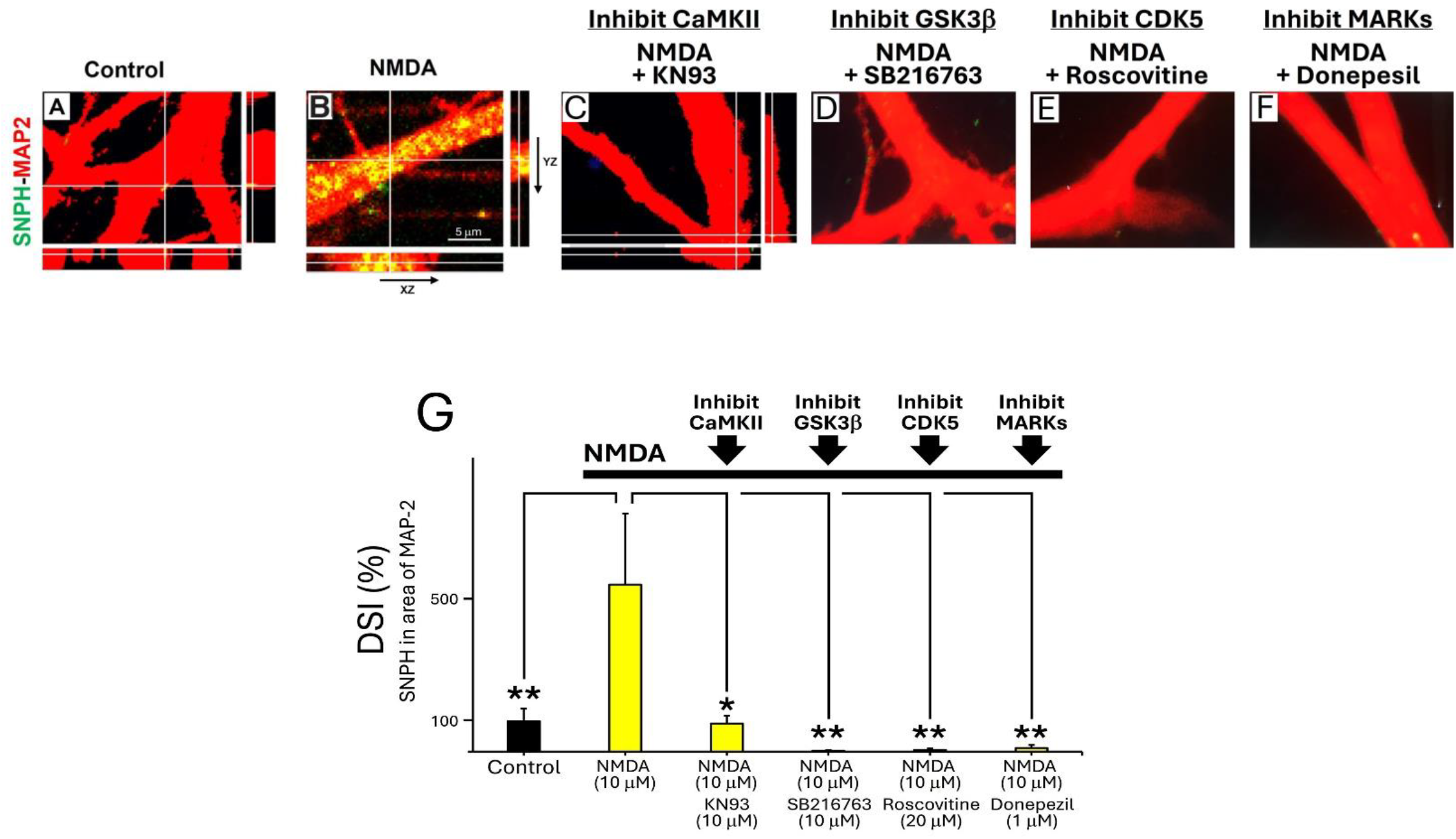
Inhibition of p-Tau kinases CAMKII, GSK-3β, CDK5, MARK prevents NMDA from triggering DSI. SNPH-MAP2 merged images showing NMDA (10 μM) triggered DSI (B) blocked by CAMKII inhibitor KN93 (10μM; C), GSK3β inhibitor SB216763 (10 μM, D), CDK5 inhibitor Roscovitine (20 μM, E) and MARKs inhibitor Donepezil (1 μM, F). G: Quantification of kinase inhibitors on DSI induced by NMDA (10μM) by pre-treating cells with 30 min of inhibitors before NMDA was applied for 24 hrs. N=dendrites for each group are N=51,63,31,30,30,30 for Control, NMDA, KN93, SB216763, Roscovitine and Donepezil respectively. One-way Anova with Post Hoc Tukey HSD (beta) test (* p<0.05, ** p<0.01). Orthogonal (xz, yz) images only shown for A, B, C.

Strikingly, inhibiting these three p-Tau kinases with inhibitors completely prevented NMDA from triggering DSI (Fig. 5 D,E,F,G). We used inhibitor concentrations based on published work (GSK-3β inhibitor SB216763 (10 μM) [25, 26]; CDK5 inhibitor Roscovitine (20 μM) [22]; MARKs inhibitor Donepezil (1 μM) [27]). Collectively, the p-Tau kinase inhibition data in Fig. 5 suggests that NMDA utilizes p-Tau as a downstream step to drive DSI.

### 3.4. Aβ42 differentially activates GluN2B to drive DSI by p-Tau

Having shown that NMDA differentially utilizes GluN2B to drive DSI by p-Tau (Fig. 3-5), we next examined if another key pathogen, namely Aβ42, also differentially activates GluN2B to drive DSI by p-Tau. The role of Aβ42 in DSI is an important issue, given Aβ42 is a key pathogen in AD and that there is comorbidity between AD and MS [2]. A 2022 published work [28] showed that in cultured neurons, extracellular Aβ42 (2.5 μM for 48 hrs) colocalizes selectively with GluN2B and not GluN2A. We therefore suggest that Aβ42 might selectively use GluN2B as a mediator to drive DSI. Excitingly, we demonstrated in Fig. 6 that Aβ42 (5 μM, 24 hrs) does indeed trigger DSI (B). Importantly, the DSI triggered by of Aβ42 is completely blocked by inhibiting GluN2B with RO-25 (C), with quantitative results shown in (D).

**Fig. 6.**
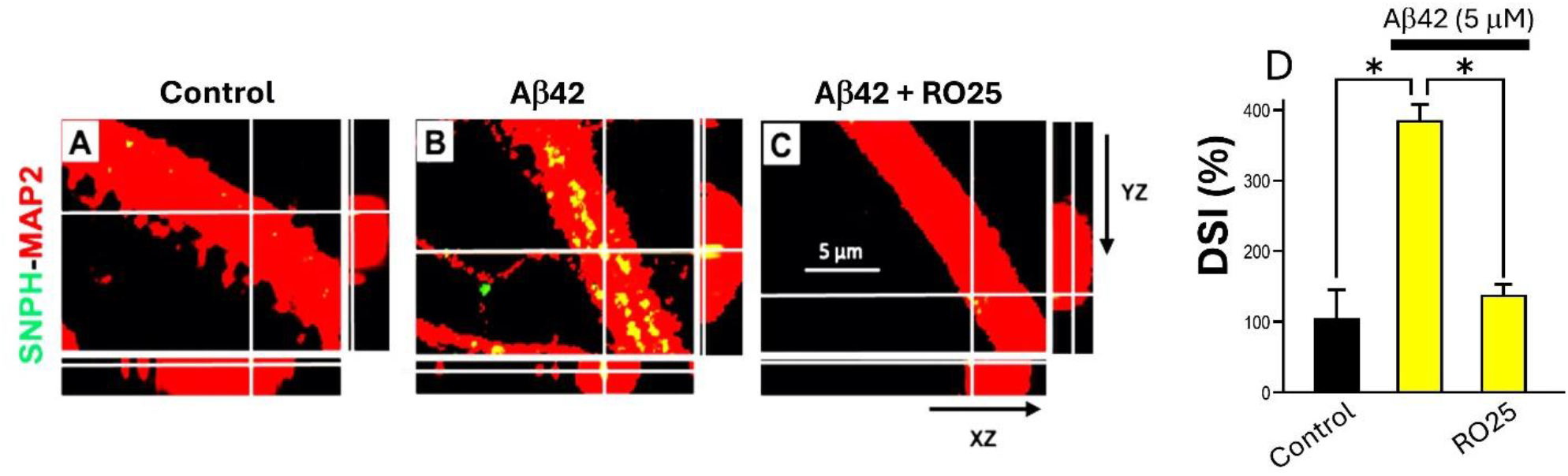
Inhibition of GluN2B completely prevents Aβ42 from driving DSI. Cells pre-treated with 30 min of RO25 (10 mM) before β-amyloid (5µM) was applied for 24 hrs. N=30 dendrites for each group. One-way Anova with Post Hoc Tukey HSD (beta) test (*p<0.05). No difference between Control vs RO25+ Aβ42 (p = 0.89)

Having shown that Aβ42 utilizes GluN2B to drive DSI, we next examined if GluN2B drives DSI using a p-Tau kinase with major relevance to AD. It well established that DAPK1 is a kinase that contributes to p-Tau in AD [29], and inhibiting DAPK1 has been shown to reduce Tau phosphorylation levels in both cell culture and mouse models of AD [30]. We therefore examined if inhibiting DAPK1 prevents DSI. Fig. 7 shows that inhibiting DAPK1 with the potent inhibitor (4Z)-4-(3-Pyridylmethylene)-2-styryl-oxazol-5-one (100 nM) [31] completely prevents NMDA from triggering DSI (Fig. 7 B, C). Quantitative results are shown in Fig. 7D.

**Fig. 7.**
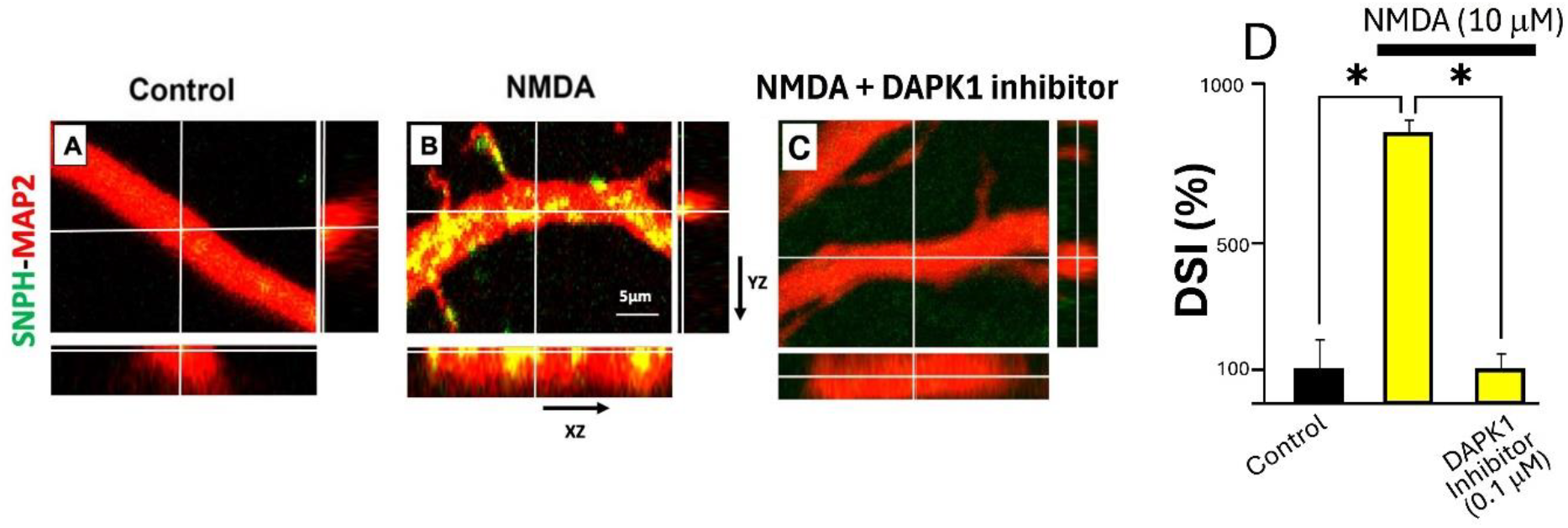
Inhibition of DAPK1 completely prevents NMDA from driving DSI. DSI triggered by NMDA (10μM) is prevented by DAPK1 inhibitor (4Z)-4-(3-Pyridylmethylene)-2-styryl-oxazol-5-one (0.1μM). One-way Anova with post-hoc Tukey HSD test was applied between the experimental conditions. Significant differences were observed between NMDA (10μM) and DAPK1 inhibitor (0.1μM); p<0.05. For control, no of dendrites = 30; for NMDA treated neurons, no of dendrites= 40; for DAPK1 inhibitor (0.1μM), no of dendrites= 30.

Collectively, our results strongly suggest that DSI driven by p-Tau underlies the comorbidity between AD and MS [2].

### 3.5. Tau-KO prevents DSI

One potential pitfall of Fig. 5 & 7 is that inhibiting various p-Tau kinases could inhibit DSI by affecting other proteins unrelated to Tau. In Fig. 8, we directly implicate Tau by Tau-KO. We cultured primary hippocampal neurons from Tau-KO mice (B6.129X1-Mapttm1Hnd/J, Jackson) then applied NMDA to assay DSI. Amazingly, Fig. 8 shows that unlike wild type (Tau +/+) neurons that generate robust DSI, Tau-/-neurons completely prevent NMDA from driving DSI (Fig. 8).

**Fig. 8.**
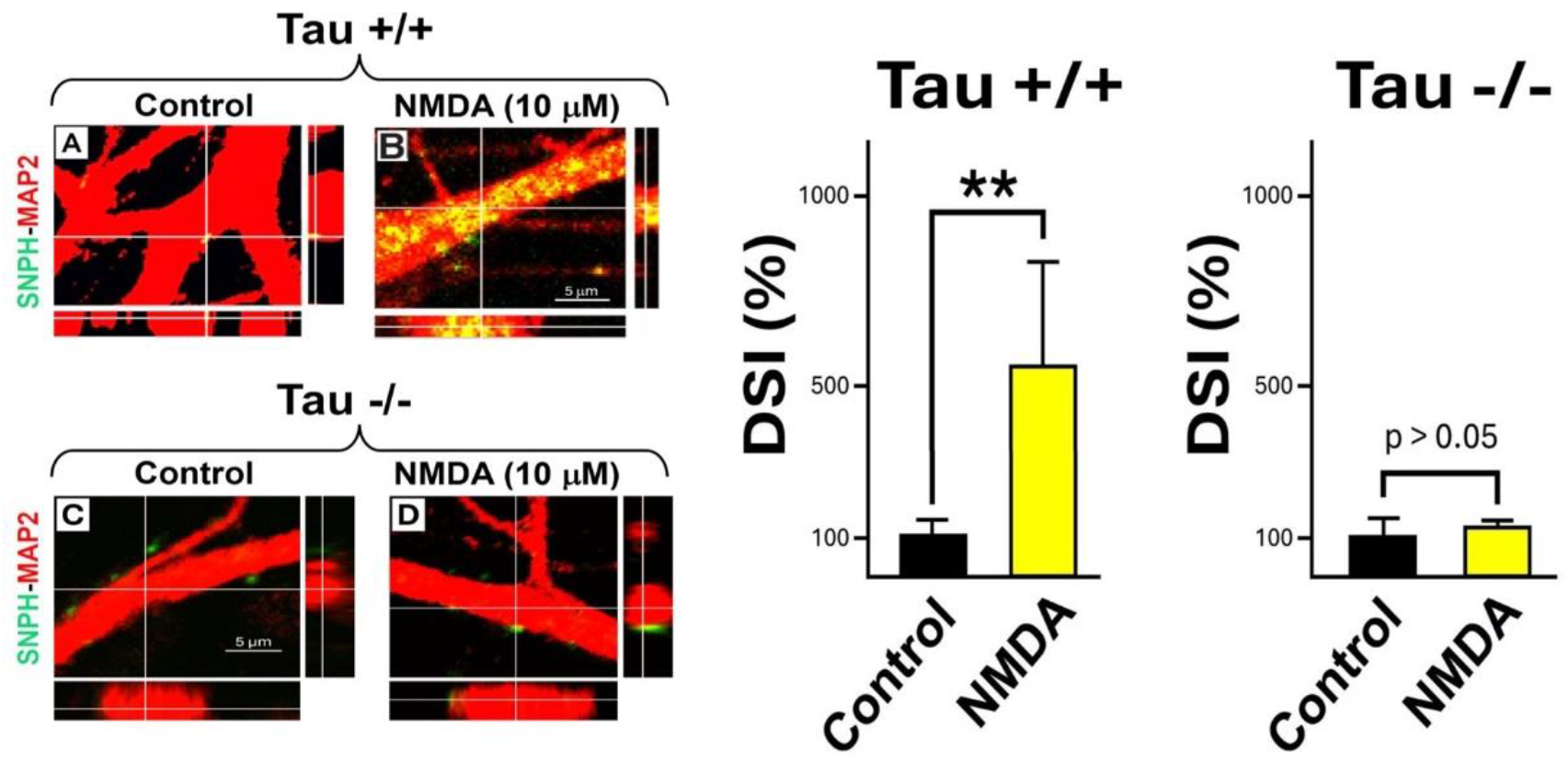
Tau-KO abolishes DSI induction by NMDA. (A, B) **<u>Tau +/+ neurons</u>**. Merged images of SNPH-MAP2 staining of cultured wild type (Tau+/+) hippocampal neurons showing NMDA (10μM, 24 hrs) triggered massive DSI (B). Quantification of DSI shown on right. N=51, 63 dendrites for Control and NMDA, One-way Anova with Post Hoc Tukey HSD (beta) test (** p<0.01). Same data taken from Fig. 5 (C, D) **<u>Tau -/- neurons</u>**. Same experiments performed in Tau-/- hippocampal neurons showed complete abolishment of DSI by Tau deletion. Orthogonal 3-D analysis showed Green SNPH puncta resided only outside, but never inside, Red MAP2 dendrites. Quantification of DSI shown on right. Three experiments with N=30 dendrites for each group, One-way Anova with Post Hoc Tukey HSD (beta) test (no significant difference, p >0.05)

Collectively, abolishing DSI using various p-Tau kinase inhibitors (Fig. 5 & 7) and direct confirmation using Tau-KO (Fig. 8) prove that Tau is an indispensable link to driving the toxic DSI.

## 4. Discussion

The key finding of this study is to reclassify MS, classically a white matter disease characterized by demyelination, into a tauopathy disease of the gray matter, particularly in the incurable Progressive MS phase. Existing studies from other studies have already established that tau phosphorylation is driving multiple sclerosis (MS) pathology [7], and in the progressive phase of MS patients, cerebrospinal fluid (CSF) p-Tau increases over time, reaching the highest levels in the late phase of primary progressive MS (PPMS) [16]. In this study, we demonstrated that tauopathy is a not a passive ‘bystander’ but actively drives a novel toxic target in models of progressive MS, the DSI, recently discovered in our laboratory [17]. The key supporting data is that inhibitors of p-Tau kinases (Fig. 5 & 7) and Tau-KO (Fig. 8) completely abolish DSI in cultured neurons.

Anderson et al. demonstrated enormous tau hyperphosphorylation in brains of primary progressive patients. In particular, there was a preponderance of insoluble, abnormally phosphorylated tau [32] along with glial cells which is a characteristic feature of neurodegenerative diseases. Interestingly, studies conducted using CSF of PPMS patients revealed diverse results. Some studies found increased levels of p-tau in PPMS compared to healthy controls but not significantly different from other MS clinical subtypes like RRMS and SPMS [23]. Others showed mild elevation of p-tau in a few MS patients during active phases [24, 26]. This indicates that upregulation of p-tau protein does not always correlate with neurological worsening [23]. CSF tau biomarkers provide some information but remain inconsistent for PPMS, so more longitudinal, PPMS-specific studies are required.

Studies in chronic experimental autoimmune encephalomyelitis (EAE), an animal model of MS, revealed the presence of excessively phosphorylated tau protein and insoluble tau assembled at different neuronal sites. The hyperphosphorylated tau protein levels were linked with neuronal damage, especially as the disease progressed to chronic or secondary progressive stage [15, 25]. Hyperphosphorylated tau accumulates and forms insoluble aggregates in brain tissue of SPMS patients associated with neuronal damage [25].

In another study conducted in MOG induced EAE model, variation of tau protein expression at different neuronal sites was observed as the disease progressed from one stage to another. In early disease course, tau protein was localized mainly in nerve fibres. However, when the disease reached the chronic stage, tau protein expression was found more in neuronal cell bodies. Levels of p-tau protein were found elevated and linked with neuroaxonal damage, whereas total tau concentrations were found decreased—suggesting that p-tau may drive long-term neurodegeneration more so than total tau [22]. Studies performed in P301S-htau transgenic mice, which overexpress human pathogenic tau demonstrated an increased expression of neuronal hyperphosphorylated tau. This was accompanied with early axonal death, gliosis and infiltration of cytokines [27].

So far, no tau-targeted therapy has been tested specifically in PPMS clinical trials. However, the presence of tauopathy in PPMS suggests potential relevance. ASO due to their specificity appear especially promising for future investigation, given their progress in Alzheimer’s disease and other tau-related pathology. Still, none have yet been explored in MS—even preclinically. Altogether, these data point towards tauopathy as a potential contributor to neurodegeneration in PPMS. Our key finding in this paper that tauopathy drives a novel toxic mitochondrial target to damage gray matter in animal models of progressive MS could lead to using Tau ASO knockdown therapy to treat progressive MS.

## 5. Conclusions

Multiple Sclerosis (MS) has a late Progressive MS phase involving neurodegeneration of gray matter with no known cure. Understanding what damages Progressive MS is critical for therapy development. We propose that Progressive MS is damaged by tauopathy driving a novel toxic mitochondrial target (DSI) into neuronal dendrites to kill gray matter by excitotoxicity. Accordingly, we propose a future therapy for Progressive MS involving using Antisense Oligonucleotides (ASO) to knock down Tau to ameliorate this toxic mitochondrial target to restore neuronal health. Excitingly, there is already an existing Tau ASO therapy approved by FDA in 2025 for clinical trials in AD. This could expedite a breakthrough treatment for Progressive MS, by repurposing Tau ASO therapy in AD to MS. We are currently initiating Tau ASO therapy in animal models of Progressive MS as a prelude to repurpose the FDA Tau therapy from AD to clinical trials in MS patients.

